# RNF43 truncating mutations mediate a tumour suppressor-to-oncogene switch to drive niche-independent self-renewal in cancer

**DOI:** 10.1101/748574

**Authors:** Maureen Spit, Nicola Fenderico, Ingrid Jordens, Tomasz Radaszkiewicz, Rik G.H. Lindeboom, Jeroen M. Bugter, Lars Ootes, Max Van Osch, Eline Janssen, Kim E. Boonekamp, Katerina Hanakova, David Potesil, Zbynek Zdrahal, Sylvia F. Boj, Jan Paul Medema, Vitezslav Bryja, Bon-Kyoung Koo, Michiel Vermeulen, Madelon M. Maurice

## Abstract

Wnt/β-catenin signalling is a primary pathway for stem cell maintenance during tissue renewal and a frequent target for mutations in cancer. Impaired Wnt receptor endocytosis due to loss of the ubiquitin ligase RNF43 gives rise to Wnt-hypersensitive tumours that are susceptible to anti-Wnt-based therapy. Contrary to this paradigm, we identify a class of RNF43 truncating cancer mutations that strongly induce β-catenin-mediated transcription, despite exhibiting retained Wnt receptor downregulating activity. Mechanistically, these RNF43 mutants trap Casein Kinase (CK)1 at the plasma membrane, which prevents β-catenin turnover and propels ligand-independent Wnt target gene transcription. When introduced in human colon stem cells, these oncogenic RNF43 mutants cooperate with p53 loss to drive a niche-independent program for self-renewal and proliferation. Importantly, onco-RNF43 mutations, unlike conventional LOF RNF43 mutations, confer resistance to anti-Wnt-based therapy. Our data demonstrate the relevance of studying patient-derived mutations for understanding disease mechanisms and improved applications of precision medicine.

---

Aberrant activation of Wnt/β-catenin signalling is a key oncogenic event that confers an undifferentiated state and allows cancer cells to thrive outside their native niche constraint^1–3^. In adult stem cells, Wnt signalling is curbed by the negative feedback regulators RNF43 and ZNRF3, two homologous transmembrane ubiquitin ligases that induce removal of the Wnt receptors FZD and LRP6 from the cell surface via ubiquitin-mediated endocytosis and lysosomal degradation^4,5^. Within the stem cell niche, the activity of RNF43/ZNRF3 is counterbalanced by secreted proteins of the R-spondin (Rspo) family that form a complex with Leucine-rich repeat-containing G-protein coupled receptor 4/5 (Lgr4/5) to mediate membrane clearance of RNF43/ZNRF3, promote Wnt receptor stabilization and enhance Wnt responsiveness of stem cell populations^6–13^.

Mutational loss of RNF43 and/or ZNRF3 is observed in human malignancies of the colon, pancreas, stomach, ovary, endometrium and liver^14–20^. Inactivation of RNF43/ZNRF3-mediated feedback leads to an increased abundance of Wnt receptors at the cell surface, which renders cells hypersensitive to Wnt ligands in their environment^5^. The resulting Wnt-dependent growth state drives tumorigenesis and generates a druggable addiction to Wnt ligands in these cancer subsets^5,15,17,18,21,22^. Indeed, blocking Wnt ligand biogenesis by small molecule inhibitors of the O-acyltransferase Porcupine (PORCN) suppresses the growth of RNF43-deleted pancreatic and small intestinal tumors in preclinical models^15,23,24^. This vulnerability has offered an opportunity to treat human cancers that are genetically defined by RNF43 mutations with Porc inhibitors. Currently, four small-molecule PORCN inhibitors are evaluated in clinical trials for cancer treatment (ClinicalTrials.gov, NCT01351103, NCT02278133, NCT02521844, NCT03447470, NCT02675946, and NCT03507998).

Clearly, mutational inactivation of RNF43 is a prerequisite for Wnt-dependent growth^23^. Recent cancer genome sequencing efforts, however, revealed a large diversity of genetic lesions within the *RNF43* locus of various human cancer types (cBioportal.org)^25^. This mutational variability poses a challenge to unambiguously distinguish driver from passenger mutations and predict which mutations truly generate a Wnt-dependent growth state that can be exploited by targeted treatment. Insight in the mechanisms by which individual RNF43 mutations contribute to cancer development and progression therefore is vital for the understanding of patient-specific disease mechanisms and the development of precision oncology strategies.

Here, we uncover a class of RNF43 truncating mutations that drive inappropriate Wnt pathway activation by a mechanism distinct from RNF43 LOF mutations. Through trapping of Casein kinase 1 (CK1) at the plasma membrane, these RNF43 mutants interfere with the turnover of the transcriptional coactivator β-catenin, promoting the transcriptional activation of Wnt target genes. When introduced in primary human colon stem cells, truncated RNF43 mutants induce a state of oncogenic stress and require prior inactivation of TP53 to drive a niche-independent program for self-renewal and proliferation. Importantly, oncogenic RNF43 mutations, unlike conventional LOF RNF43 mutations, confer resistance to anti-Wnt-based therapy. Our results reveal the functional heterogeneity of cancer driver mutations in a single gene and demonstrate the importance of examining patient-derived mutations to uncover disease mechanisms, allow for improved patient stratification and applications of targeted therapy.

## Results

### Loss of the C-terminus endows the tumor suppressor RNF43 with oncogenic properties

RNF43 comprises a single-span transmembrane E3 ubiquitin ligase of 783 amino acids (Fig. 1a). Binding and ubiquitination of Wnt receptors maps to the N-terminal half of the RNF43 protein, including the extracellular (ECD), transmembrane (TM) and RING domains. These domains are followed by an extended C-terminal tail that contains conserved Ser-, His-, and Pro-rich regions to which no role has been assigned (Fig. 1a). Notably, a third of reported *RNF43* cancer variants comprise nonsense or frameshift mutations that prospectively yield expression of C-terminally shortened RNF43 proteins for which functional consequences remain unknown (cBioportal.org)^20,25^. To address this issue, we expressed RNF43 cancer variants carrying incremental C-terminal truncations in HEK293T cells and monitored their impact on Wnt-induced β-catenin-mediated transcription. Strikingly, RNF43 variants truncated between residues K514-Q563 strongly induced β-catenin-mediated transcription, independent of supplementation with Wnt (Fig. 1b). Unlike a well-defined LOF missense variant (I48T)^26^, these RNF43 mutants fully retained their ability to downregulate FZD receptors while strongly increasing cytosolic β-catenin levels (Fig. 1c-f). Wnt pathway activation by the representative truncated RNF43 variant R519X remained unaffected by PORCN inhibitors^24^, while these compounds fully suppressed Wnt signalling activity induced by the LOF cancer mutant I48T or by *RNF43/ZNRF3* deletion (Fig. 1g, Supplementary Fig. 1a)^15^. Thus, in contrast to LOF mutations, RNF43 R519X induces pathway activation in a ligand-independent manner. Furthermore, RNF43 R519X strongly induced Wnt pathway activation in an *RNF43/ZNRF3*-knockout background, indicating that these mutants do not operate via a dominant-negative mechanism (Supplementary Fig. 1a). Thus, truncated RNF43 variants gain competence to drive basal Wnt signalling and are functionally distinct from classical LOF mutations.

**Figure 1.**
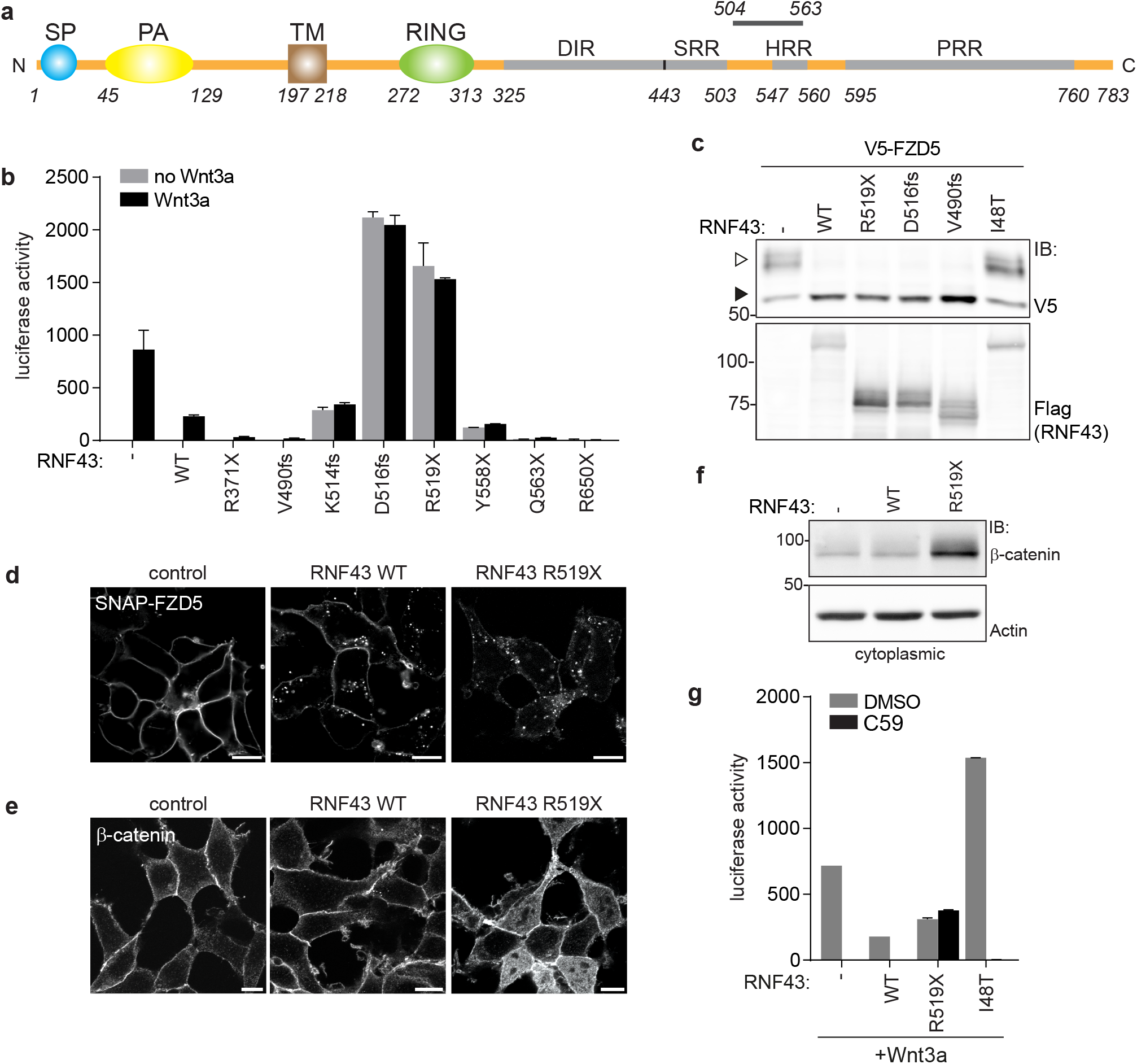
Cancer truncations endow RNF43 with oncogenic properties. **a,** Schematic representation of the RNF43 protein. SP; signal peptide, PA; protease-associated domain, TM; transmembrane domain, RING; E3 ligase catalytic domain, DIR; Dishevelled-interaction region, SRR; Serine-rich region, HRR; Histidine-rich region, PRR; Proline-rich region. The region in which oncogenic truncations occur (aa 504-563) is indicated. **b,** Wnt luciferase-reporter activity in HEK293T cells expressing the indicated RNF43 cancer mutants. Average luciferase reporter activities ±s.d. in n = 2 independent wells are shown. **c,** Western blot analysis showing the effect of RNF43 cancer mutants on V5-FZD5 expression in HEK293T cells. Open and closed arrows indicate mature (post-Golgi) and immature (ER-associated) FZD5, respectively. **d,** Confocal microscopy of surface labeled SNAP-FZD5 in HEK293T cells upon expression of RNF43 WT or R519X. Cells were chased for 30 min. Scale bars represent 10 μm. **e,** Confocal microscopy of β-catenin localisation in HEK293T cells expressing RNF43 WT or R519X. Scale bars represent 10 μm. **f,** Western blot analysis of RNF43 WT and R519X for cytoplasmic β-catenin levels. **g,** Wnt luciferase-reporter activity in HEK293T cells expressing Wnt3a and WT RNF43, oncogenic RNF43 (R519X) or a LOF RNF43 variant (I48T) after treatment with DMSO or the PORCN inhibitor C59. IB; immunoblot, WT; wild-type.

### Premature termination codons within the oncogenic region of RNF43 avoid nonsense-mediated decay

More precise mapping using designed RNF43 truncations showed that truncations within a region ranging from D504-Q563 unleash β-catenin-mediated transcription, indicating that oncogenic activity requires retention of the Ser-rich region and loss of the His- and Pro-rich regions (Fig 1b, Supplementary Fig. 1b). Mutations introducing premature termination codons (PTC) within this *RNF43* region occurred in various cancer types, including pancreas, endometrium, ovarium and colon (Supplementary Table 1). Expression of inappropriately truncated proteins is commonly limited due to nonsense-mediated decay mRNA surveillance pathways^27,28^. To investigate this issue, we employed CRISPR/Cas9 to introduce biallelic *RNF43* PTCs in SW480 APC-mutant colorectal cancer cells, in which *RNF43* is actively transcribed (Supplementary Fig. 1c, d). Mutant *RNF43* mRNAs (V520fs/D516fs) were expressed even at increased abundance compared to parental cells (Supplementary Fig. 1e, f), indicating that these transcripts are stable and give rise to a protein product that promotes transcription via a feedforward mechanism.

### Truncated RNF43 cancer variants interfere with downstream Wnt signalling events

Next, we aimed to identify the molecular requirements for the oncogenic activity of truncated RNF43 variants. Conventional wild-type RNF43 tumour suppressor activity relies on the RING-type E3 ligase domain that marks FZD for ubiquitin-mediated endocytosis and lysosomal turnover (Fig. 1c, d)^4,5^. The introduction of RING domain-inactivating mutations still allowed for RNF43 R519X-mediated induction of basal β-catenin transcription, while responses to Wnt were further enhanced, likely due to the FZD stabilising effects of this catalytically inactive RNF43 variant (Fig. 2b)^5^. Furthermore, RNF43 R519X retained its ability to drive basal Wnt pathway activation when the ECD and TM domains were substituted by those of the unrelated transmembrane proteins CD16 and CD7 (Fig. 2a, c). Similar replacements in full-length RNF43 resulted in failure to downregulate FZD5 and loss of Wnt inhibitory activity (Fig. 2c)^29^. Thus, the truncated RNF43 cytosolic tail is required and sufficient to drive oncogenic β-catenin-dependent transcription by a mechanism independent of the ECD and RING domains. In line with this, RNF43 R519X-induced β-catenin transcription was insensitive to expression of Dishevelled (Dvl)-1 DEP-C, a Dvl fragment that binds the FZD cytosolic domains and blocks Wnt-mediated receptor activity (Fig. 2d)^30^. In addition, signalling induced by ectopic expression of β-catenin was unaffected by RNF43 R519X (Fig. 2e). We conclude that truncated RNF43 cancer variants affect a molecular step positioned downstream of the Wnt receptors and upstream of β-catenin-mediated transcription.

**Figure 2.**
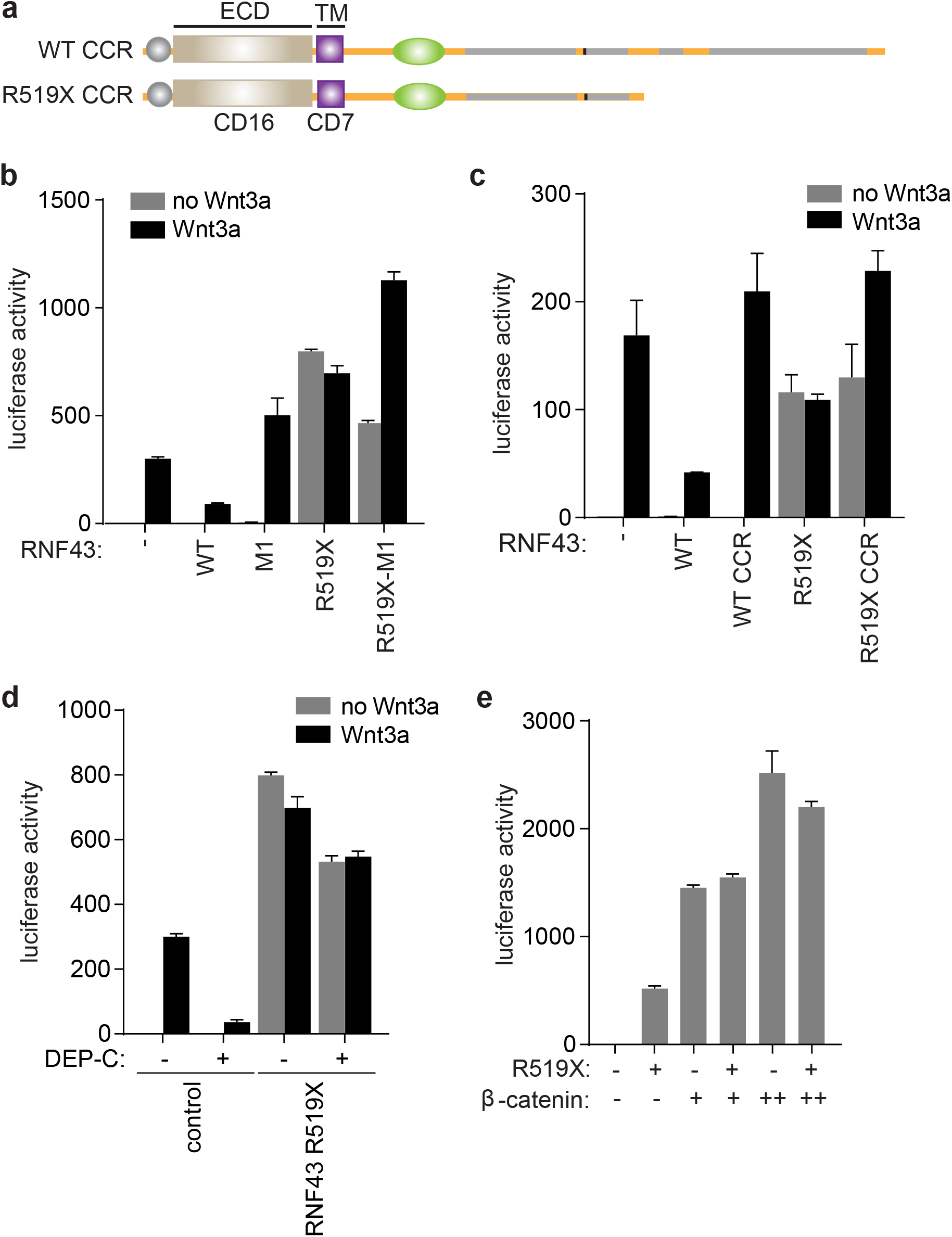
Oncogenic RNF43 variants activate Wnt signalling downstream of the receptor complex and upstream of β-catenin. **a,** Schematic of RNF43 constructs in which the extracellular and transmembrane domains (ECD and TM) are replaced by those of the unrelated CD16 and CD7, respectively. CCR; CD16-CD7-RNF43. **b,** Wnt luciferase-reporter activity in HEK293T cells expressing WT RNF43 and oncogenic RNF43 (R519X) harbouring mutations C290S/H292S in the RING domain (M1). Average luciferase reporter activities ±s.d. in n = 2 independent wells are shown. **c,** Wnt luciferase-reporter activity in HEK293T cells expressing the indicated RNF43 constructs. Average luciferase reporter activities ±s.d. in n = 2 independent wells are shown. **d,** Wnt luciferase-reporter activity in HEK293T cells co-expressing oncogenic RNF43 (R519X) and the Dishevelled DEP-C tail. Average luciferase reporter activities ±s.d. in n = 2 independent wells are shown. **e,** Wnt luciferase-reporter activity in HEK293T cells co-expressing oncogenic RNF43 (R519X) and increasing amounts of β-catenin. Average luciferase reporter activities ±s.d. in n = 2 independent wells are shown. WT; wild-type.

### Truncated RNF43 variants retain CK1 at the plasma membrane to drive β-catenin-mediated transcription

We next investigated if truncated RNF43 interferes with the β-catenin destruction complex, which provides a central point for Wnt pathway regulation and is commonly targeted by inactivating mutations in cancer^31,32^. BioID proximity labeling revealed interactions of WT RNF43 with destruction complex members Axin1, CK1α, CK1ε and APC (Fig. 3a, Supplementary Table 2). Strikingly, interactions of truncated oncogenic RNF43 variants with endogenous Axin1, and CK1α/ε were increased in comparison to RNF43 WT (Fig. 3b, Supplementary Fig. 2a), while interactions with APC and GSK3β were not noticeably altered or even decreased (Fig. 3b, Supplementary Fig. 2b, c). Moreover, expression of RNF43 R519X, but not WT or the non-oncogenic R371X variant, prompted a redistribution of Axin1, as well as endogenous CK1α and CK1ε, from cytosol to the plasma membrane (Fig. 3c, d, Supplementary Fig. 2d, e). The interaction of CK1α/ε and RNF43 remained unaffected by depletion of Axin1 and its close homologue Axin2 (Supplementary Fig. 2f), suggesting that CK1 interacts directly with the RNF43 cytosolic tail. Binding of CK1α/ε mapped to an intermittent region in the RNF43 C-terminus, composed of S486-Q488 and G492-S494 (Supplementary Fig. 2g, h). Moreover, levels of CK1 binding correlated with Wnt pathway-activating ability of RNF43 R519X (Supplementary Fig. 2h, i). These findings imply that retention of CK1 by RNF43 truncated cancer variants is essential for their oncogenic mode of action.

**Figure 3.**
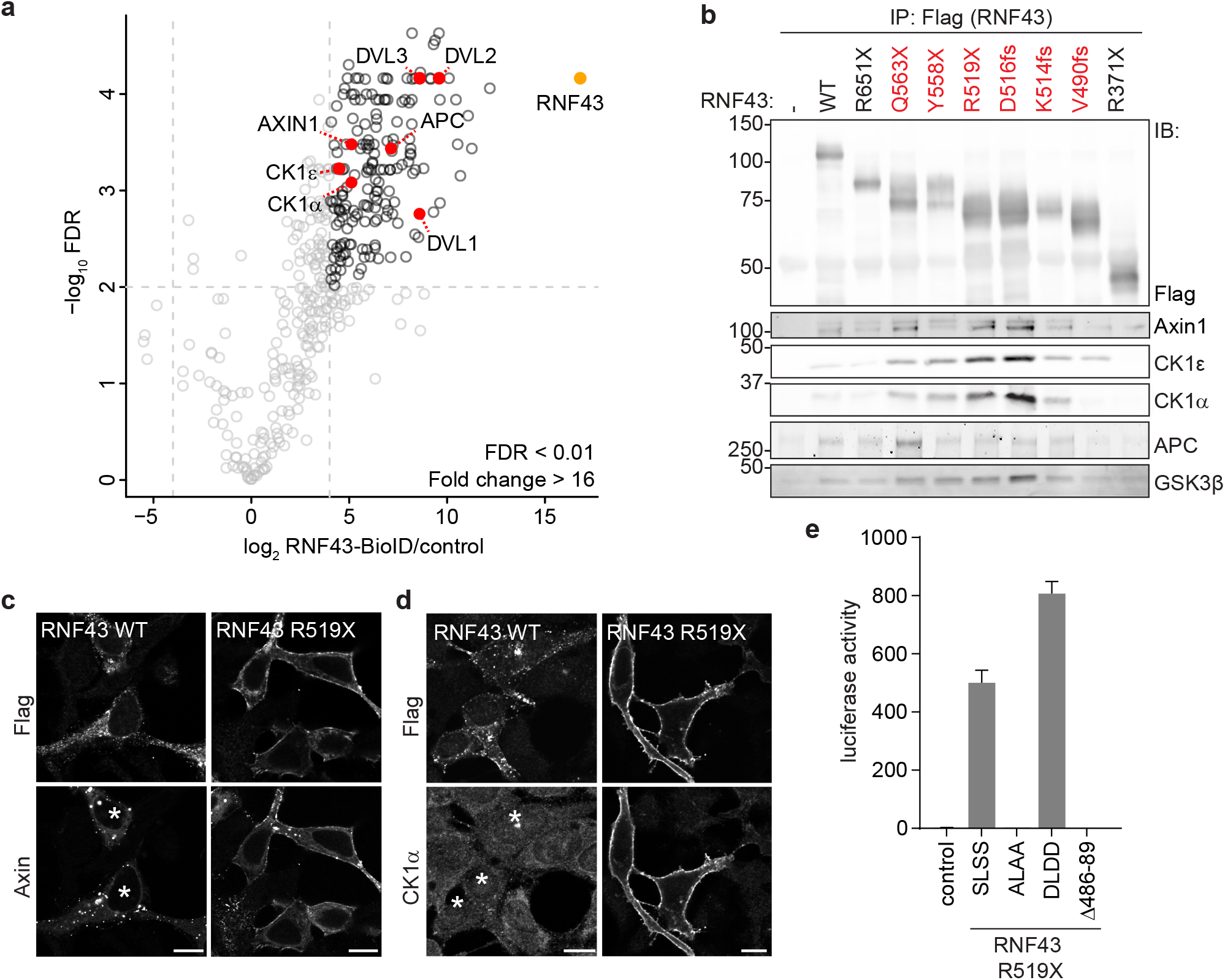
Oncogenic RNF43 variants trap components of the β-catenin destruction complex at the plasma membrane. **a,** Volcano plot showing proteins enriched after streptavidin pull-down of biotin-treated HEK293 cells stably expressing dox-inducible RNF43-BirA*. Striped line demarcates the empirical 0.01 False discovery rate (FDR) cut off. Significantly enriched Wnt/β-catenin pathway components are highlighted in red. **b,** Western blot analysis of endogenous destruction complex components that co-precipitated with the indicated RNF43 cancer variants expressed in HEK293T cells. Oncogenic RNF43 truncations are indicated in red. **c,d** Confocal microscopy analysis of the subcellular localisation of Axin1-GFP **(c)** and endogenous CK1α **(d)** upon expression of WT or oncogenic RNF43 (R519X). Scale bars represent 10 μm. RNF43 is visualised by Flag-staining. Asterisks indicate RNF43-expressing cells. **e,** Wnt luciferase-reporter activity in HEK293T cells expressing non-mutated RNF43 R519X (SSLS) or the ALAA, DLDD and Δ486-89 mutants. Average luciferase reporter activities ±s.d. in n = 2 independent wells are shown. IP; immunoprecipitation, IB; immunoblot, WT; wild-type.

### CK1-mediated phosphorylation of the RNF43 cytosolic tail confers oncogenic Wnt pathway activation

To investigate whether RNF43 is a target for CK1-mediated Ser/Thr phosphorylation, we analysed the phosphorylation status of RNF43 WT and R519X. Indeed, phosphorylation of RNF43 R519X was markedly increased when compared to RNF43 WT, consistent with its ability to trap endogenous CK1 (Supplementary Table 3). Phosphorylation of WT RNF43 was increased upon co-expression of CK1α, but not CK1ε, suggesting that CK1α is the preferred kinase for functional cooperation. We identified a non-canonical CK1 SLSS target sequence at residues 500-503, which when truncated, abolishes oncogenic activity of RNF43 (Supplementary Fig. 1b). Ala substitution of the SLSS motif (SLSS>ALAA) in full-length RNF43 abolished pathway suppression, while introduction of phospho-mimetic residues (SLSS>DLDD) promoted suppressor activity, indicating that WT RNF43 normally employs CK1 kinase activity to perform its tumour suppressor role (Supplementary Fig. 2j). In accordance, an RNF43 cancer variant carrying a CK1 binding site deletion (ΔS486-G489>R; cBioportal) displayed LOF effects (Supplementary Fig. 2j). Introduction of SLSS>ALAA or CK1 binding site deletion fully abrogated the capacity of RNF43 R519X to induce basal Wnt pathway activation, while SLSS>DLDD R519X mediated increased tumourigenic activity (Fig. 3e). Thus, truncated RNF43 employs CK1 binding and phosphorylation to drive oncogenic Wnt pathway activation.

### Oncogenic RNF43 mutations induce a TP53-dependent growth arrest in human colon organoids

Next, we investigated the impact of oncogenic RNF43 truncations on epithelial homeostasis, using human colon organoids^33,34^. Introduction of CRISPR/Cas9-mediated frame shift mutations within the endogenous *RNF43* locus (onco-RNF43) yielded only a limited number of small organoid clones that failed to thrive, reminiscent of a senescent phenotype (Supplementary Fig. 1c, 3a)^35^. Genotyping of a slowly expanding clone confirmed the presence of a mono-allelic onco-RNF43 mutation (Supplementary Fig. 3b). This phenotype is strikingly different from *RNF43* LOF mutations that are well tolerated in intestinal organoids^5,36^. We wondered how onco-RNF43 induces epithelial growth arrest. In line with our model of onco-RNF43-mediated CK1 sequestration, ablation of *Csnk1a1* (CK1α) from the mouse intestinal epithelium was shown previously to trigger massive Wnt pathway activation accompanied with p53-mediated cellular senescence^37^. Combined ablation of *Csnk1a1* and *Tp53* instigated formation of highly invasive carcinomas^37^. Similarly, we noted a co-occurrence of oncogenic *RNF43* frameshift mutations with mutations in *TP53* or senescence-associated genes in human cancer (Supplementary Table 1), suggesting that *TP53* inactivation might be required to bypass an oncogenic stress-induced growth arrest. Indeed, combined onco-*RNF43*/*TP53*KO mutant clones rapidly appeared after CRISPR/Cas9 targeting, thrived in large numbers and allowed for the occurrence of bi-allelic onco-*RNF43* mutations (Supplementary Fig. 3a, b). Thus, loss of TP53 creates a permissive cellular state for onco-RNF43 expression.

### Onco-RNF43 variants drive niche-independent growth in human colon organoids and confer resistance to anti-Wnt-based therapy

A key feature of cancer pathway driver mutations is their ability to confer niche-independent growth, which is examined by depleting stem cell growth factors from the organoid culture medium^1,34^. We wondered if the ability of onco-RNF43 to drive basal β-catenin-mediated transcription alleviates the need for supplementation of colon organoids with Wnt and/or R-spondin (Rspo), a Wnt-potentiating niche factor that induces membrane clearance of RNF43/ZNRF3 and allows for enhanced Wnt responsiveness of stem cell populations^6–8,10–12,38^. Omitting Wnt readily compromised viability of both WT and *TP53*KO organoid lines, as reported earlier^34,39^, while onco-*RNF43/TP53*KO organoid growth remained unaffected (Fig. 4a). Although instant removal of Rspo was not tolerated by any of the organoid lines, onco-*RNF43*/*TP53*KO organoids displayed much greater tolerance to a step-wise decrease in Rspo concentrations when compared to WT and *TP53*KO organoid lines (Fig. 4a). We conclude that onco-RNF43 mutations confer decreased dependence on Wnt and Rspo niche factors, a hallmark of cancer cell growth.

**Figure 4.**
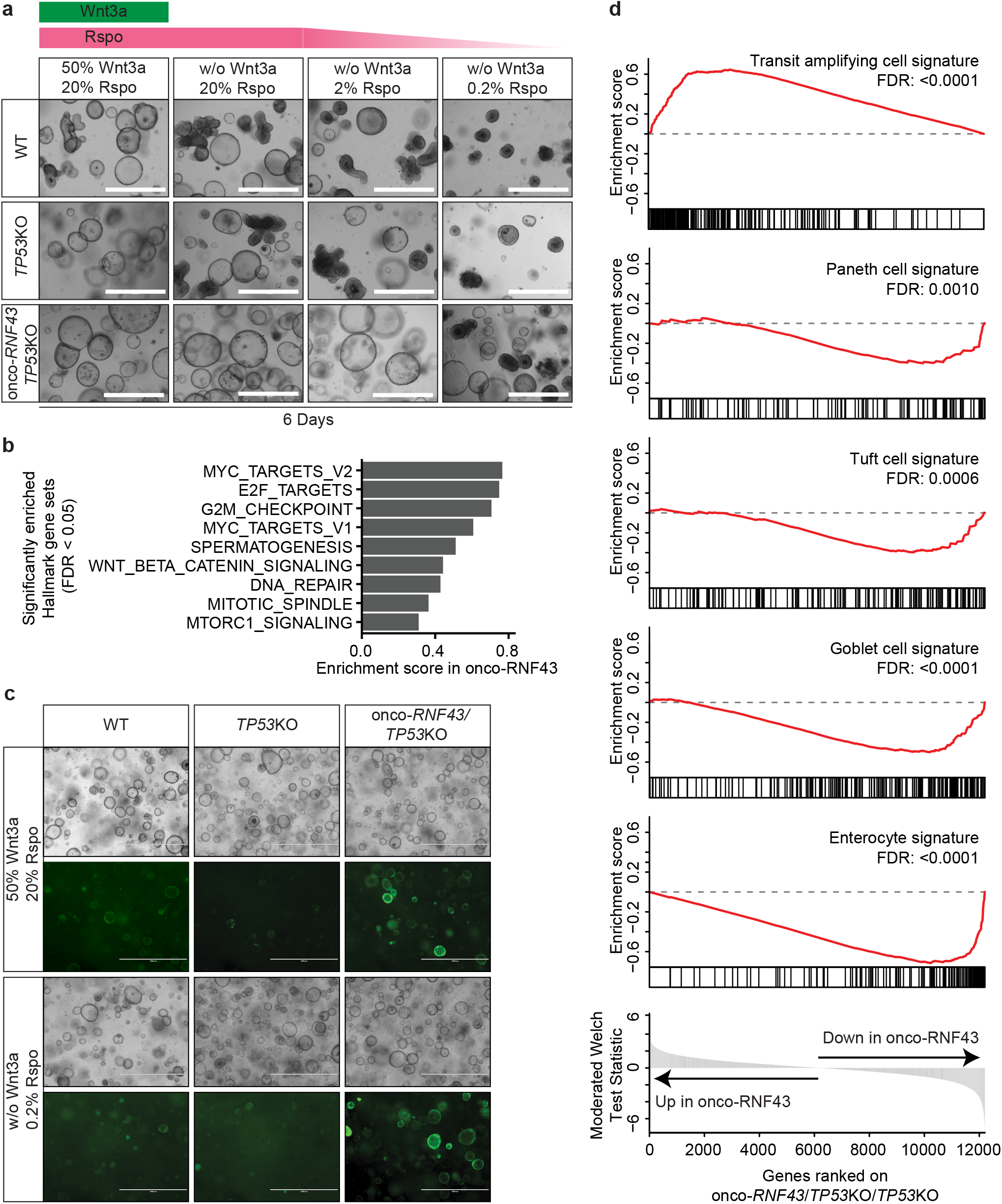
Onco-*RNF43/TP53* KO organoids display an oncogenic transcriptional profile that drives self-renewal and niche-independent growth. **a,** Bright-field microscopy images of WT, *TP53*KO and onco-*RNF43/TP53*KO human colon organoid lines grown in medium with high Wnt/Rspo (20% conditioned medium (CM)) or without Wnt/Rspo (20, 2 or 0.2% CM). Scale bars represent 1000 μm. **b,** Bar plot showing the enrichment scores of significantly enriched MSigDB hallmark gene sets in onco-*RNF43/TP53*KO compared to *TP53*KO organoids (FDR < 0.05). **c,** Bright-field microscopy and fluorescence microscopy pictures of WT, *TP53*KO and onco-*RNF43/TP53*KO human colon organoid lines grown in two different media and transduced with the TOP-GFP reporter. Scale bars represent 1000 μm. **d,** Gene Set Enrichment Analysis of onco*-RNF43/P53*KO compared to *TP53*KO organoids in medium without Wnt/low Rspo (0.2%). Significantly changed intestinal cell-type gene sets from Haber et al.^42^ are shown (FDR < 0.05).

To investigate the impact of onco-RNF43 mutations on gene expression in colon epithelial cells, we performed RNA sequencing of WT, *TP53KO* and onco-*RNF43/TP53KO* organoid lines grown in high Wnt/Rspo (20%) or no Wnt/low Rspo (0.2%) medium. Unsupervised clustering of significantly changing genes revealed four distinct clusters of gene expression dynamics (Supplementary Fig. 4a). Onco-RNF43-mediated transcriptome alterations were markedly enhanced in no Wnt/low Rspo growth conditions (Supplementary Fig. 4a, b). Therefore, we focused on 1448 genes that were differentially expressed in *TP53*KO versus onco-*RNF43/TP53*KO organoid lines grown in no Wnt/low Rspo (Supplementary Fig. 4a, b). Of the 966 downregulated genes in onco-*RNF43*/*TP53*KO organoids, a minor group of 123 genes was traced back to loss of *TP53*. The remaining 843 genes were specifically and consistently downregulated by onco-RNF43 expression. Conversely, 482 genes were significantly increased in onco-RNF43-expressing organoids. Noticeably, this gene set was also upregulated in high Wnt/Rspo-treated organoids (Supplementary Fig. 4a, right heatmap), indicating that onco-RNF43 confers a signature normally provided by the stem cell niche. Gene set enrichment analysis (GSEA) using MSigDB^40^ revealed significant enrichment for Myc and E2F targets, Wnt signalling, DNA damage response and cell division by onco-RNF43 expression (Fig. 4b). A lentiviral Wnt GFP reporter^41^ confirmed sustained Wnt signalling in onco-RNF43-expressing organoids in no Wnt/low Rspo growth conditions (Fig. 4c). Furthermore, differentiated cell type signatures^42^ were lost in onco-RNF43-expressing organoids while profiles of transit amplifying cells were notably enriched (Fig. 4d). In summary, onco-RNF43 induces expression of a stem cell-like transcriptome in colonic epithelial cells and these effects are intensified in conditions where Wnt and Rspo are scarce.

The ability of onco-RNF43 mutations to drive Wnt-*independent* signalling stands in stark contrast to the previously described role of RNF43 LOF mutations that promote a Wnt-*dependent* growth state. Importantly, our findings predict differential sensitivity of these RNF43 mutational classes to treatment with PORCN inhibitors, Wnt antagonists that are currently evaluated for clinical treatment of RNF43-mutant cancer patients^3,43–45^. Indeed, a large fraction of onco-*RNF43*/*TP53*KO organoid clones survived and recovered after one week of treatment with PORCN inhibitor C59, while no viable clones were obtained for WT and *TP53*KO organoids (Fig. 5a, b). Thus, onco-RNF43 expression confers resistance of human colon organoids to PORCN inhibitor treatment.

**Figure 5.**
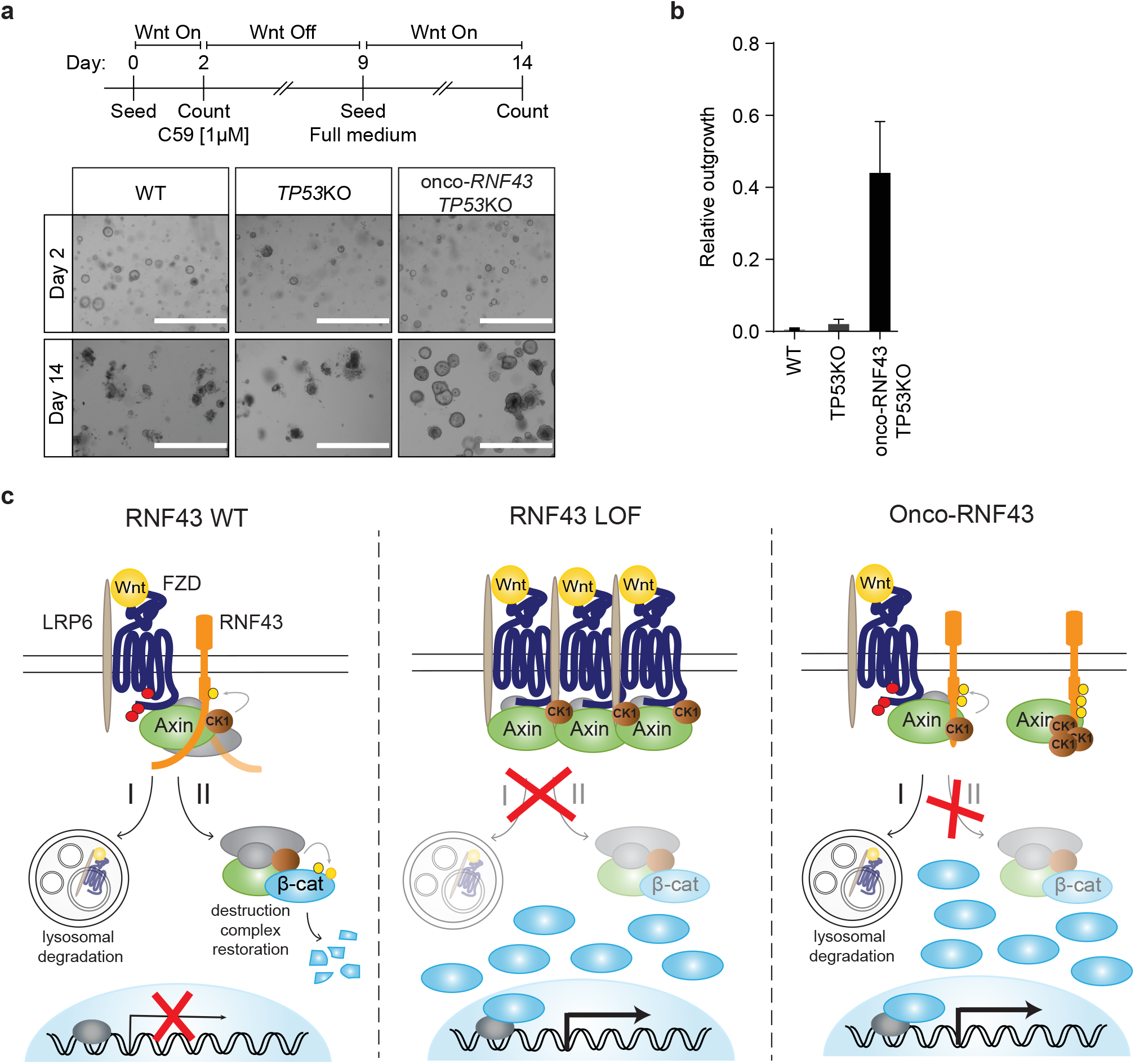
Onco-*RNF43/TP53* KO colon organoids are resistant to Porcupine inhibitors. **a,** Top: Schematic experimental set up of the clonogenic assay. Bottom: Bright-field microscopy images of WT, *TP53*KO and onco-*RNF43/TP53*KO human colon organoid lines at day 2 and 14. Scale bars represent 1000 μm. **b,** Relative outgrowth of WT, *TP53*KO and onco-*RNF43/TP53*KO organoid lines treated with the PORCN inhibitor C59 (1 μM) for 7 days. Graph shows the outgrowth of organoids at 5 days after splitting (day 14) relative to organoids at day 2. Error bars represent ±s.d. of the mean of n = 3 experiments. **c,** Model for mode of action RNF43 LOF and onco-RNF43 variants. (Left) RNF43 performs a bifunctional tumour suppressor role by (I) targeting Wnt receptors for endocytosis and lysosomal degradation, and (II) by transiently interacting with the destruction complex to reconstitute its activity in the cytosol and re-establish Wnt pathway inhibition. This second suppressor role involves CK1-mediated phosphorylation and an unknown molecular activity of the flexible cytosolic tail of RNF43. (Middle) Classical loss-of-function (LOF) mutations prevent RNF43 function at the plasma membrane, leading to Wnt receptor overexpression and, consequently, hypersensitivity of cancer cells to Wnt. (Right) Onco-RNF43 truncated variants are stably expressed at the plasma membrane and retain E3 ligase-dependent Wnt receptor downregulating activity. However, due to loss of the cytosolic tail, these mutants selectively trap CK1 and Axin1 at the plasma membrane, which prevents destruction complex assembly and drives uncontrolled β-catenin-mediated transcription of target genes. WT; wild-type, LOF; loss-of-function, β-cat; β-catenin.

## Discussion

Inappropriate Wnt/β-catenin signalling in cancer is achieved by two major mutational driver routes. Inactivation of the destruction complex is well-studied and exemplified by the prominent driver role of APC mutations in colorectal cancer^46^. More recently, misregulation of Wnt receptor abundance emerged as an alternative oncogenic pathway^3,47^. RNF43 mutations, found in ~19% of colorectal cancer cases, are mutually exclusive to APC mutations and considered a prime hallmark of Wnt-hypersensitive cancer subsets^15,20,23^. Our study uncovers a class of RNF43 mutations that drive inappropriate Wnt pathway activation by a mechanism distinct from that of RNF43 LOF or APC mutations. By trapping CK1 at the membrane, these RNF43 mutants compromise β-catenin destruction complex activity in the cytosol, leading to stabilisation of β-catenin and the transcriptional activation of Wnt target genes (Fig. 5c). In support of their oncogenic role, the activity of these RNF43 variants does not require inactivation of the paralogue ZNRF3^5^. Different from APC mutations^48^, onco-RNF43 mutations induce a state of oncogenic stress in primary cells and require prior inactivation of TP53 to exert their oncogenic role.

In summary, we identified a class of RNF43 mutations that mediate a tumour suppressor-to-oncogene switch to drive downstream Wnt pathway activation. Unlike LOF mutations, the expression of onco-RNF43 variants predicts prolonged survival in Wnt-depleted conditions and thus provides a contraindication for Wnt- or receptor-inhibiting treatment strategies. Our findings further imply that WT RNF43 performs a bifunctional tumour suppressor role, mediating ubiquitin-dependent Wnt receptor downregulation^4,5^ as well as ubiquitin-independent regulation of destruction complex activity (Fig. 5c). Our study demonstrates the importance of examining patient-derived mutations to identify novel tumourigenic molecular mechanisms, obtain a broader comprehension of signalling pathways in normal and cancer cells and improve applications of precision cancer medicine.

## METHODS

### Cell culture and transfection

Human Embryonic Kidney (HEK) 293 cells, HEK293T cells and SW480 were cultured in RPMI or DMEM high glucose (Invitrogen), respectively, supplemented with 10% fetal bovine serum (GE Healthcare), 2 mM UltraGlutamine (Lonza), 100 units/mL penicillin and 100 μg/mL streptomycin (Invitrogen). Cells were cultured at 37 °C in 5% CO_2_. Wnt3a-conditioned medium (CM) was obtained from L-cells stably expressing and secreting Wnt3a^49^ cultured in DMEM low glucose (Invitrogen). Rspo- and Noggin-CM were produced as described before^50^. For luciferase reporter assays, HEK293T cells were stimulated overnight (o/n) and for protein expression experiments cells were stimulated for 3 h with Wnt3a-CM. Transfections were performed using FuGENE 6 (Promega) according to manufacturer’s protocol. siRNA transfection was performed using Lipofectamine™ RNAiMAX (Thermo Fisher) according to manufacturer’s protocol at 50nM for 4 days. Control (#1) siRNA were obtained from Ambion (Thermo Fisher) and the Axin1 SMARTpool of siRNA from Dharmacon (Horizon). For co-transfection, plasmids were transfected 1 day before analysis.

### Plasmids and antibodies

Flag-Dishevelled1 DEP-C, TOPFlash and FOPFlash luciferase reporter plasmids were described previously^30,49^. Myc-β-catenin and mouse Wnt3a were a kind gift of Hans Clevers (Hubrecht Institute, Utrecht, Netherlands). RNF43–2×Flag–HA and RING mutants were described previously^5^. Plasmid for expression of human Axin1-GFP was described previously^51^. All mutants were generated by either site-directed mutagenesis or by PCR-subcloning using Q5 High-Fidelity 2x Master Mix (NEB). All constructs were sequence verified. The following primary antibodies were used for immunoblotting (IB), immunofluorescence (IF) or immunoprecipitation (IP): goat anti-Axin1 (R&D systems), goat anti-CK1ε (Santa Cruz), goat anti-CK1α (Santa Cruz), rabbit anti-GSK3β (Cell Signaling), mouse anti-β-catenin (BD Transduction laboratories), rabbit anti-APC (Santa Cruz), rabbit anti-FLAG (Sigma-Aldrich), rat anti-HA (Roche), rabbit anti-V5 (Sigma-Aldrich), mouse anti-FLAG (M2; Sigma-Aldrich), mouse anti-V5 (Genscript), mouse anti-GFP (Roche), mouse anti-Actin (MP Biomedicals). Primary antibodies were diluted conform manufacturer’s instructions. Secondary antibodies used for IB or IF were used 1:8000 or 1:300 respectively and obtained from either Rockland or Invitrogen.

### TOPFlash luciferase reporter assays

HEK293T cells were seeded in 24-well plates and transfected the next day with 30 ng of reporter construct TOPFlash or FOPFlash, 5 ng of thymidine kinase (TK)-Renilla and the indicated constructs. Cells were stimulated 6 h post-transfection with Wnt3a-CM o/n, then cells were lysed in Passive lysis buffer (Promega) for 20 min at room temperature (RT). IWP-2 (R&D systems) was used o/n at 5 μM. Levels of Firefly and Renilla luciferase were measured using the Dual Luciferase Kit (Promega) accordingly to the manufacturer’s instructions on a Berthold luminometer Centro LB960.

### Cell lysis and immunoprecipitation

HEK293T were transfected with polyethylenimine (PEI). 24 hours post-transfection, cells were washed and collected in ice-cold PBS, and subsequently lysed in lysis buffer (50 mM Tris, pH 7.5, 150 mM NaCl, 0.5% Triton X-100, 5 mM EDTA, 1 mM DTT, 50 mM sodium fluoride and protease inhibitors) for 30 min on ice followed by 30 min centrifugation at 4 °C. Supernatants were used for immunoprecipitations (IPs) using 1 μg of the indicated antibody. Samples were left tumbling for 1 h at 4 °C, followed by 1 h incubation with protein A or G beads (RepliGen and Millipore, respectively). For FLAG IPs, 15 μL pre-coupled FLAG M2 agarose (Sigma-Aldrich) was added for 1.5 h at 4 °C while tumbling. Agarose beads were washed six times with lysis buffer and proteins were eluted in SDS sample buffer by 5 min boiling, or for FZD samples, incubated for 45 min at 37 °C.

### Cell fractionation

24 hours post-transfection, HEK293T cells were washed and collected in PBS and subsequently incubated for 10 min on ice in fractionation buffer (10 mM HEPES pH 7.9, 1.5 mM MgCL2, 10 mM KCl, 1mM DTT and protease inhibitors) to allow the cells to swell. Cells were homogenized by 25-50 strokes in a Douncer after which the homogenization was visually evaluated by microscopy. The homogenate was centrifuged at 500 × g at 4 °C for 10 min to obtain the nuclear pellet. The supernatant was subsequently centrifuged at 100.000 × g at 4 °C for 1 h to separate the membrane fraction from the cytosolic fraction.

### Immunoblotting

Western blotting was performed using standard procedures with Immobilon-FL PVDF membranes (Millipore). In short: after protein transfer, the membranes were blocked for 1 h at RT in 1:1 ratio Odyssey blocking buffer (LI-COR): PBS. Primary antibodies were incubated o/n at 4 °C and secondary antibodies for 1 h at RT in the dark. The LI-COR Odyssey infrared imaging system was used for immunoblot analysis.

### Immunofluorescence and SNAP labeling

HEK293T cells were grown on laminin-coated glass coverslips in 24-well plates and transfected after 24 h. For SNAP labeling, cells were labelled with 1 mM SNAP-surface^549^ (Bioke) for 15 min at RT, washed and subsequently chased for 30 min at 37 °C. Cells were fixed in 4% paraformaldehyde or ice-cold methanol and blocked in PBS containing 2% BSA and 0.1% saponin. Primary and secondary antibody incubations were performed in blocking buffer for 45 min – 1 h at RT. Cells were mounted in ProLong Gold (Life Technologies) and analysed using Zeiss LSM510 or LSM700 confocal microscope.

### BioID for RNF43 interacting proteins

For the identification of RNF43 binding proteins a previously described protocol was used^52^. Briefly, pcDNA4-TO-RNF43-BirA*-HA was obtained by cloning RNF43 into pcDNA3.1 MCS-BirA(R118G)-HA (a gift from Kyle Roux; Addgene # 36047), which was subcloned into pcDNA4-TO. pcDNA4-TO-RNF43-BirA*-HA was transfected into T-REx™-293 cells and selected with 200 μg/ml of zeocin to obtain a stable RNF43 TetON -REx™-293 BioID cell line. Ten days after transfection, single cells were plated in 96-wells and grown in the presence of 100 μg/ml zeocin and 5 μg/ml of blasticidin. For validation the selected clones were analyzed by Western blot and immunofluorescence for tetracycline (1 μg/mL, Santa Cruz Biotechnology) induced expression of RNF43-BirA*-HA fusion protein and enzyme activity of BirA* by biotin supplementation (50 μM, Santa Cruz Biotechnology). To identify RNF43 interacting proteins, TetON -REx™-293 BioID cells were seeded in 15 cm dishes and were treated o/n with tetracycline and biotin. Non-induced cells were used as a negative control. Cells were lysed in 2 mL of lysis buffer containing 2% TX-100, 500 mM NaCl, 0.2% SDS, 50 mM Tris, pH 7.4 supplemented with protease inhibitors (Roche), phosphatase inhibitors (Calbiochem), 1 mM DTT and PIC (Roche). Lysates were collected, sonicated and cleared by centrifugation at 16.500 × g for 15 min at 4 °C. Streptavidin beads (Streptavidin Sepharose High Performance, GE Healthcare) were added to the lysates and incubated for 16 h tumbling at 4 °C. Subsequently, beads were washed four times with lysis buffer and two times with 50 mM Tris HCl pH 7.4. Next, beads were washed with ammonium bicarbonate buffer and proteins were reduced with DTT and alkylated with iodacetamide. Trypsin digestion on beads was performed o/n at 37 °C. Resulting peptides were transferred into LC-MS vials and concentrated to 15 μL. LC-MS/MS analysis was performed on an RSLCnano coupled to an Orbitrap-Elite system. MS/MS data processing was performed using Proteome Discoverer (version 1.4). Hits are classified as proteins with a minimum of two unique peptides present in at least two out of three replicates. Proteins were filtered using the cRAP contaminant database and proteins interacting with the BirA* tag or present in the negative control were subtracted. For volcano plot analysis, only proteins identified in at least all three replicates of either the control or the BioID sample were considered. By using Perseus 1.5.5.3^53^ with default settings, missing values were imputed from a semi-random normal distribution around the lower detection limit of all detected proteins. False discovery rates (FDR) are calculated by a modified t-test (in Perseus) followed by Benjamini-Hochberg FDR adjustment.

### Identification of phosphorylation sites by mass spectrometry

Briefly, cells were seeded in 15 cm dishes and transfected at 80% of confluency with the indicated plasmids using PEI. Per dish, cells were lysed in 2 ml of lysis buffer containing 1% NP-40, 150 mM NaCl, 50 mM Tris, pH 7.5 supplemented with protease inhibitors (Roche), phosphatase inhibitors (Calbiochem), 1 mM DTT and 10 mM NEM (N-ethylmaleimide) (Sigma-Aldrich). Then lysates were collected, sonicated and cleared by centrifugation at 16.100 × g for 20 min at 4 °C. Next, 2 μg HA-11 antibody (MMS-101R, Covance) was added and samples were incubated for 1 h tumbling at 4 °C. Then 45 μL of equilibrated G protein sepharose beads (GE Healthcare,) was added to the sample and incubated 16 h tumbling at 4 °C. Subsequently, beads were washed six times with lysis buffer, mixed with 50 μL of 2X Laemmli buffer, boiled for 5 min and loaded on 8% SDS-PAGE gels and separated. Gels were fixed with 50% methanol, 10% acetic acid, stained with 0.1% Coomassie brilliant blue (Sigma-Aldrich) in 20% methanol, 10% acetic acid for 2 hours and destained using fixation solution. Next, corresponding 1-D bands were excised and processed for mass spectrometry analysis. Protein in gel pieces were alkylated, digested by trypsin and subsequently cleaved by chymotrypsin. Digested peptides were extracted from gels. 1/10 of the peptide mixture was directly analysed and the rest of the sample was used for TiO2 phosphopeptide enrichment. Both peptide mixtures were separately analysed on LC-MS/MS system (RSLCnano connected to Orbitrap Elite; Thermo Fisher Scientific). MS data were acquired in a data-dependent strategy selecting up to top 10 precursors based on precursor abundance in the survey scan (350-2000 m/z). High resolution HCD MS/MS spectra were acquired in Orbitrap analyzer. The analysis of the mass spectrometric RAW data files was carried out using the Proteome Discoverer software (ThermoFisher Scientific; version 1.4) with in-house Mascot (Matrixscience, London, UK; version 2.4.1) search engine utilisation. The phosphoRS feature was used for phosphorylation localisation and manually confirmed. Peptides with Mascot score > 20, rank 1 and with at least 6 amino acids were considered. Quantitative information assessment was done in Skyline software. Normalisation of the data was performed using the set of phosphopeptide standards (added to the sample prior phosphoenrichment step; MS PhosphoMix 1, 2, 3 Light, Sigma) and by nonphosphorylated peptides identified in direct analyses. Clusters were analysed by Orbitrap Script 2.0.

### smRNA FISH

SW480 cells were grown on coverslips for 24 h. For smFISH, samples were prepared as previously described^54^. Briefly, cells were fixed for 10 min with 4% Formaldehyde solution (Sigma-Aldrich) and 70% Ethanol o/n. Samples were then hybridized with Quasar 670 labelled RNF43 probes (Stellaris, Biosearch Technologies) and mounted to microscopy slides using Prolong Diamond Antifade (Invitrogen). Images were acquired using a deconvolution system (DeltaVision RT; Applied Precision) using 60x lens.

### gRNAs and genotyping

The pSpCas9(BB)-2A-Puro was obtained from Addgene (48139). gRNAs were generated as previously described^55^. gRNA: Onco-RNF43- AGGCTGCATGTCCACTCGCT or TAGGGCTGCAGTACACTAGG; RNF43 KO- ATTGCACAGGTACAGCGGGT; ZNRF3 KO- GCCAAGCGAGCAGTACAGCG; TP53KO-GGCAGCTACGGTTTCCGTCT (a gift from Jarno Drost, PMC, Utrecht); Axin2- GCTTCCGTGAGGATGCCCCG For genotyping, genomic DNA was isolated using QIAamp DNA micro kit (Qiagen). Primers for PCR amplification using GoTaq Flexi DNA polymerase (Promega) were as follows: *RNF43*_Fw 5’-AGTGGATCTGGAGAAAGCTA-3’, *RNF43*_Rev 5’-ATTCAGCTGTAGTCTCCTCT-3’; *TP53*_Fw 5’-CAGGAAGCCAAAGGGTGAAGA-3’, *P53*_Rev 5’-CCCATCTACAGTCCCCCTTG-3’; *AXIN2_Fw* 5’-AGCTTTCCTTCCTCCGGTCTTC-3’, *AXIN2_Rev* 5’- GGTCACTACAGACTTTGGGGCT-3’. Products were cloned into the pGEM-T Easy vector system I (Promega) and subsequently sequenced using the T7 sequencing primer.

### Organoid culture

Healthy human colon tissue was isolated to establish human intestinal organoids for a previous study^48^. Normal human colon organoids were cultured in advanced DMEM/F12 medium (Invitrogen), supplemented with B27 (Invitrogen), Nicotinamide (Sigma-Aldrich), N-acetylcysteine (Sigma-Aldrich), EGF (Peprotech), TGF-β type I receptor inhibitor A83-01 (Tocris), P38 inhibitor SB202190 (Sigma-Aldrich), Wnt3a-CM (50%), Noggin-CM (10%), Rspo1-CM (20%) (ENRW-Nic-A-S medium). Mutant TP53 organoids were cultured in the presence of 5 μM Nutlin-3 (Cayman Chemical). Mutant RNF43 organoids were initially selected by withdrawing Wnt3a-CM and Rspo1-CM. Where indicated, the percentages of Wnt3a-CM and Rspo1-CM were adjusted. All experimentation using human organoids described herein was approved by the ethical committee at University Medical Center Utrecht (UMCU; TcBio #12-093). Informed consent for tissue collection, generation, storage, and use of the organoids was obtained from the patients at UMCU.

### Organoid electroporation

Organoid electroporation was performed as previously described^56^. Briefly, organoids were grown in 10 μM Y-27632 (Selleck chemicals) and 5 μM CHIR 99021 (Tocris) without Wnt-conditioned media for 2 d before electroporation. 24 h before electroporation, 1.25% DMSO was added to the culture medium. For electroporation the NEPA21 electroporator was used with the configuration reported by Fujii *et al*.^56^. 10 μg of pSpCas9(BB)-2A-Puro (PX459-#48139) was used to generate mutant lines. Organoids were recovered for 1 d by adding medium supplemented with 1.25% DMSO, 10 μM Y-27632 and 5 μM CHIR 99021 followed by another day with 10 μM Y-27632 and 5 μM CHIR 99021. 5 d post-electroporation organoids were grown in ENRW-Nic-A-S. For CRISPR-engineered organoids, single clones were established by manual picking of individual organoids derived from single cells and genotyped. PiggyBAC overexpressing organoids were selected using 50 μg/μL Hygromycin (Merck).To visualise Wnt activity, organoids were transduced with 7xTcf-eGFP:SV40-PuroR (Top-GFP) (Addgene #24305)^41^ as described^57^ and selected with Puromycin (Invivogen; 2 μg/mL).

### Clonogenic assay

Clonogenic assay was performed as previously described^58^. Briefly, WT, *TP53*KO and onco-*RNF43/TP53*KO organoid lines were mechanically dissociated for 5 minutes by using a narrow Pasteur pipette. At day 2 organoids were counted and C59 (Tocris; 1 μM) was added for 7 days. At day 9 organoids were split and seeded in full medium and counted at day 14 (n = 3). Pictures were taken at day 2 and 14 using an Evos microscope.

### RNA Sequencing

Organoids were grown in full medium or medium without Wnt and 0.2% Rspo medium for 48 h and were lysed in RLT lysis buffer (Qiagen). RNA was obtained using the Qiagen QiaSymphony SP (Qiagen) according to the manufacturer’s protocol. Single-end reads of 75 bp were aligned to GRCh37 with STAR^59^ with parameters outSJfilterIntronMaxVsReadN, chimJunctionOverhangMin and chimSegmentMin set to 10000000, 15 and 15, respectively. Gene expression was quantified with edgeR with the human Ensembl transcript database version 74. Differential gene expression analysis was performed with DESeq2^60^, and its ‘rlogTransformation’ function was used for variance stabilisation of gene expression data. Clustering and generation of heatmaps was done with ComplexHeatmap^61^. UpSet plots were created with the UpSetR R package^62^. Intestinal cell-type specific gene sets for Gene Set Enrichment Analysis (GSEA; http://www.broad.mit.edu/gsea/) were defined as the 250 most specific cell-type signature genes per cell-type from Extended Figure Table 3 of Haber *et al*.^42^. Human orthologues of each cell-type gene set were determined with the biomaRt R package^63^. To calculate the false discovery rate of every GSEA, we performed 10,000 gene set perturbations.

### Data availability

The data that support the findings of this study are available from the corresponding author upon reasonable request. The RNA-seq data are publicly available at the NCBI GEO repository (accession number GSE129288).

## ACKNOWLEDGEMENTS

We thank members of the laboratory of M.M.M. for discussions and suggestions.Cara Jamieson for generating the R/Z dKO line and Manja Omerzu and Lars Kemp for the Axin2KO line; Jarno Drost (Princess Máxima Centre for Paediatric Oncology, Utrecht, Netherlands) for TP53 gRNA. This work is part of the Oncode Institute, which is partly financed by the Dutch Cancer Society. This work was supported by European Research Council Starting Grant 242958, the Netherlands Organization for Scientific Research NWO VICI Grant 91815604, ECHO Grant 711.013.012 and TOP Grant 91218050 (to M.M.M.), European Union Grant FP7 Marie Curie ITN 608180 “WntsApp” (to M.M.M.). V.B. was supported by the Czech Science Foundation (project no. GA17-16680S and GX19-28347X) and the Ministry of Education, Youth and Sports of the Czech Republic (MEYS CR, the project CEITEC 2020 (LQ1601)). CIISB research infrastructure project LM2015043 funded by MEYS CR is acknowledged for the financial support of the LC-MS/MS measurements at the CEITEC Proteomics Core Facility. M.V. is supported by an ERC Consolidator Grant (771059).

## AUTHOR CONTRIBUTIONS

M.S., N.F., I.J., and M.M.M. conceived and designed the experiments. M.S., N.F., I.J., T.R., J.M.B., L.O., M.v.O., E.J., K.H., D.P. and Z.Z. performed the experiments. M.S., N.F., I.J., T.R., R.G.H.L., J.B., L.O., M.v.O., E.J. J.P.M., V.B., B.K.K., M.V. and M.M.M analysed the data. S.F.B. provided essential reagents. M.S., N.F., I.J., T.R., R.G.H.L., V.B., M.V. and M.M.M wrote the manuscript, which was reviewed by all authors.

## Supplementary Figure legends

**Supplementary Figure 1 | Onco-RNF43 induces cytosolic accumulation of β-catenin and its endogenous mRNA transcripts are stably expressed. a,** Wnt luciferase-reporter activity in HEK293T double knock out (dKO) RNF43/ZNRF3 (R/Z) cells expressing RNF43 WT and oncogenic RNF43 (R519X) after treatment with DMSO or the PORCN inhibitor IWP-2 (o/n). Average luciferase reporter activities ±s.d. in n = 2 independent wells are shown. **b,** Wnt luciferase-reporter activity in HEK293T cells expressing the indicated RNF43 truncations in the absence of Wnt3a. Average luciferase reporter activities ±s.d. in n = 2 independent wells are shown. **c,** Schematic representation of the targeted exon of human RNF43. Oncogenic region is indicated in red. **d,** Sanger sequencing of the PCR amplification products of the mutated RNF43 alleles in SW480 cells. **e,** Fluorescence images of smFISH showing individual RNF43 mRNA dots in WT SW480 cells and cells carrying mutated RNF43 alleles. DAPI (blue) is used for nuclear staining. Scale bar represents 10 μm. **f,** Graph indicating the mean number of mRNAs for RNF43 per cell for the indicated conditions. IB; immunoblot, WT; wild-type.

**Supplementary Figure 2 | Onco-RNF43 binds and relocates the destruction complex proteins Axin1, CK1α and CK1ε to the plasma membrane. a,** Western blot analysis of RNF43 WT and oncogenic RNF43 (R519X) co-precipitated with endogenous Axin1 in HEK293T cells. **b,** Western blot analysis of RNF43 WT and oncogenic RNF43 (P524X) co-precipitated with V5-APC. **c,** Western blot analysis of myc-GSK3β co-precipitated with RNF43 WT and oncogenic RNF43 (P524X). **d,** Confocal microscopy analysis of subcellular endogenous CK1α localisation upon expression of RNF43 WT, oncogenic RNF43 (R519X) and R371X. RNF43 proteins are visualised by Flag staining. Asterisks indicate RNF43-expressing cells. Scale bars represent 10 μm. **e,** Confocal microscopy analysis of the subcellular endogenous CK1ε localisation upon expression of RNF43 WT and oncogenic RNF43 (R519X). RNF43 proteins are visualised by Flag staining. Asterisks indicate RNF43-expressing cells. Scale bars represent 10 μm. **f,** Western blot analysis of endogenous Axin1, CK1ε and CK1α co-precipitated with RNF43 WT and oncogenic RNF43 (R519X) expressed in Axin2KO HEK293T cells treated with control or Axin1 siRNA for 4 days. **g,** Western blot analysis of endogenous CK1ε co-precipitated with the indicated RNF43 variants expressed in HEK293T cells. **h,** Mapping the CK1 interaction region in oncogenic RNF43 (R519X) using triple alanine scanning. Western blot analysis of endogenous CK1α, CK1ε and Axin1 that co-precipitated with the indicated oncogenic RNF43 alanine mutants in HEK293T cells. Asterisk indicates bleed through of RNF43 WT. **i,** Wnt luciferase-reporter activity in HEK293T cells induced by oncogenic RNF43 (R519X) triple alanine mutants. Average luciferase reporter activities ±s.d. are shown. **j,** Wnt luciferase-reporter activity in HEK293T cells expressing full length RNF43 WT, ALAA, DLDD or Δ486-89 mutants. Average luciferase reporter activities ±s.d. in n = 2 independent wells are shown. IP; immunoprecipitation, IB; immunoblot, WT; wild-type.

**Supplementary Figure 3 | Ablation of *TP53* is permissive for growth of onco-***RNF43***-expressing organoids. a,** Strategy to engineer the indicated human colon organoid lines. Bright field microscopy pictures of WT, onco-*RNF43, TP53KO* and onco-*RNF43*/*TP53KO* human colon organoids. Scale bars represent 400 μm. **b,** Sanger sequencing confirmation of onco-RNF43, TP53KO and onco-RNF43/TP53KO mutations in human colon organoids.

**Supplementary Figure 4 | Comparison of onco-*RNF43* induced transcriptional alterations in human colon organoids grown in high Wnt/high Rspo and no Wnt/low Rspo conditions. a,** Bright-field microscopy pictures of WT, mono-allelic onco-*RNF43*, *TP53*KO and onco-*RNF43/TP53*KO human colon organoids. Scale bars represent 400 μm. **b,** Heatmap showing gene expression dynamics for genes that are significantly changed between onco-*RNF43/TP53*KO and *TP53*KO (FDR < 0.01). Relative changes in gene expression are shown as row Z-scores. The first heatmap shows the dynamics between samples grown in no Wnt/low Rspo (0.2%) medium. Rows in the second heatmap are matched to the first heatmap and show gene expression dynamics in organoids grown in full medium. The right heatmap (green/purple) shows the log2 fold change between all samples in full medium compared to all samples in no Wnt/low Rspo (0.2%). Hierarchical clustering and k-means clustering was performed on row Z-scores in the first heatmap and used to identify four clusters of genes with distinct expression dynamics between the organoid lines. **c,** UpSet plot showing the number of significantly changing genes in different comparisons. Pairwise comparisons between WT, *TP53*KO and onco-*RNF43/TP53*KO organoids were performed on growth medium with high Wnt/Rspo (20%) (blue) and without Wnt/low Rspo (0.2%) (green). In addition, all samples in high Wnt/Rspo were compared to all samples in no Wnt/low Rspo medium (black). Number of significantly changing genes (FDR < 0.01) in each comparison is shown in the horizontal bar plot. Intersections of genes that change significantly in more than one comparison are shown as connected dots in the UpSet plot. The number of genes in each intersection is shown in the vertical bar plot. Only the 50 biggest intersection groups are shown. Multiple testing correction was done with the Benjamini–Hochberg procedure on all pairwise comparisons combined.

