## Supplementary figures and images for "RNF43 truncating mutations mediate a tumour suppressor-to-oncogene switch to drive niche-independent self-renewal in cancer"

### Supplementary Figure 1

#### Supplementary Figure 1

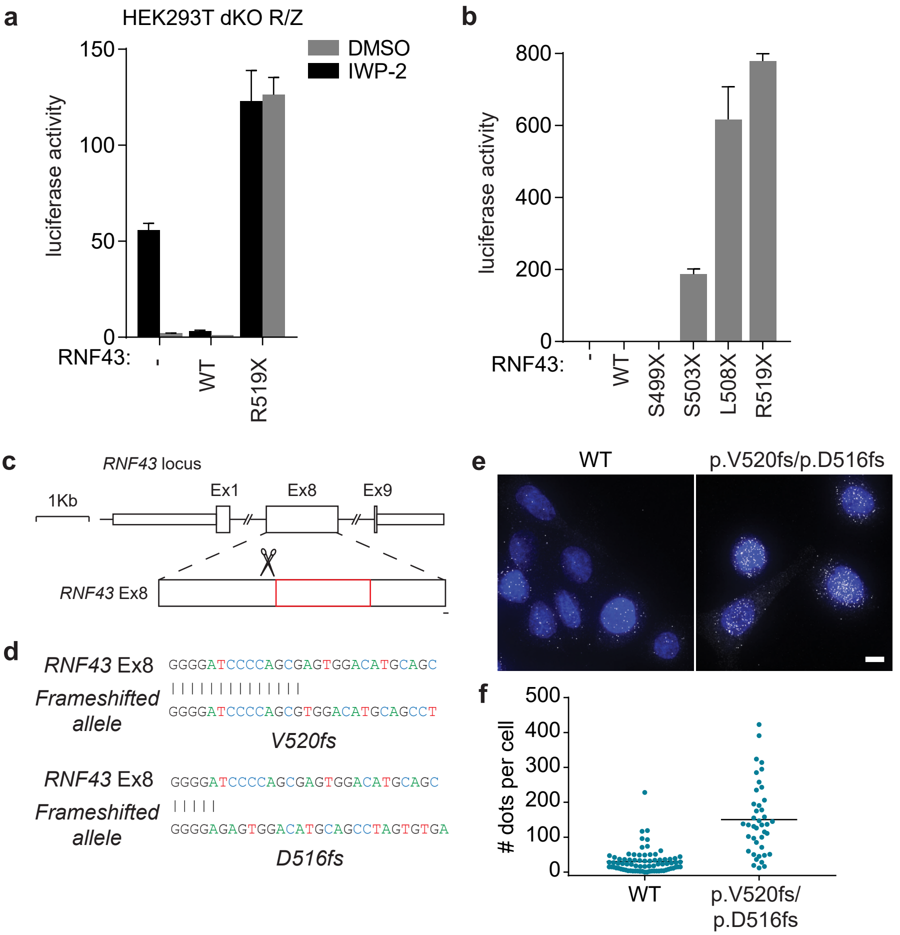

### Supplementary Figure 2

#### Supplementary Figure 2

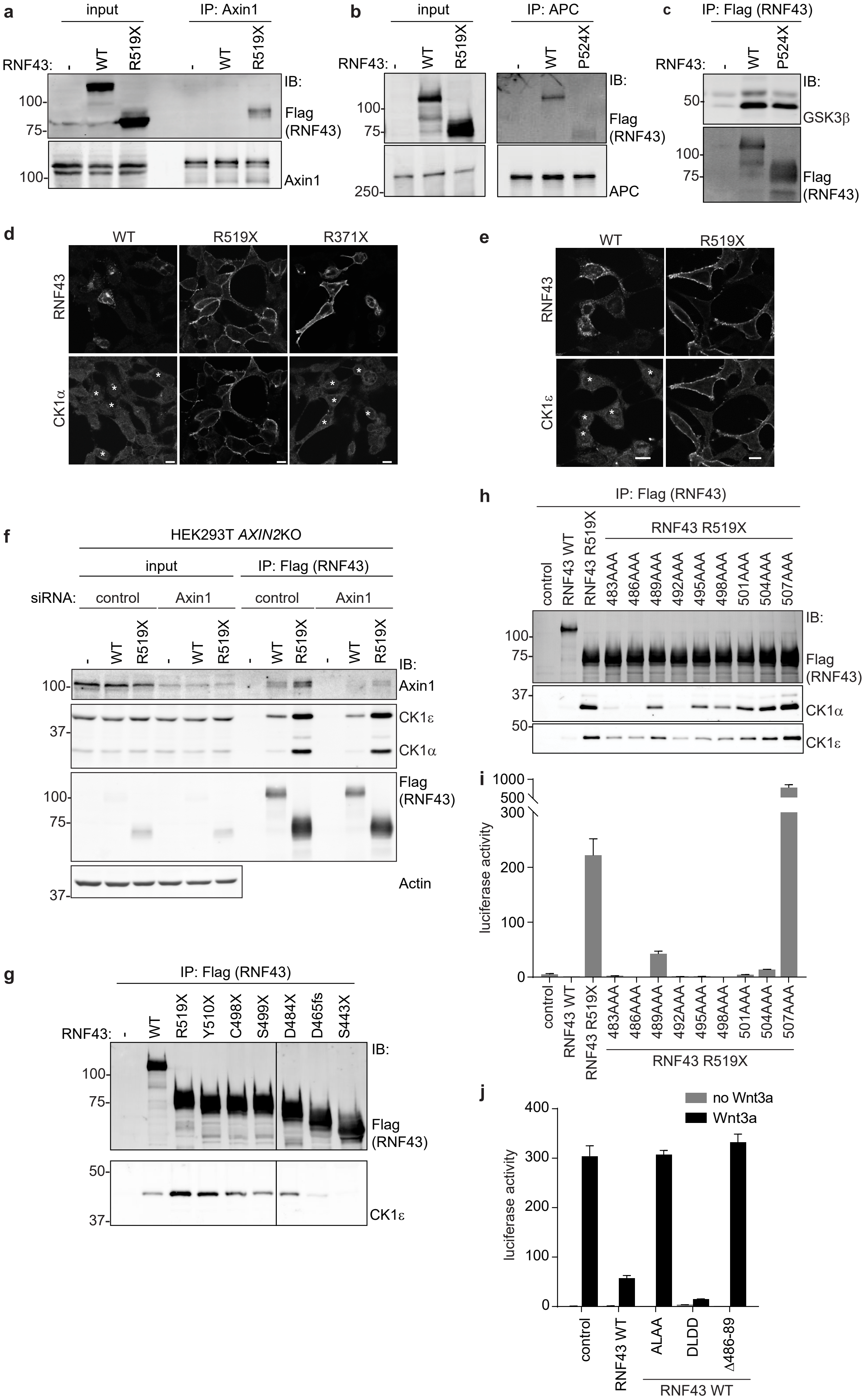

### Supplementary Figure 3

#### Supplementary Figure 3

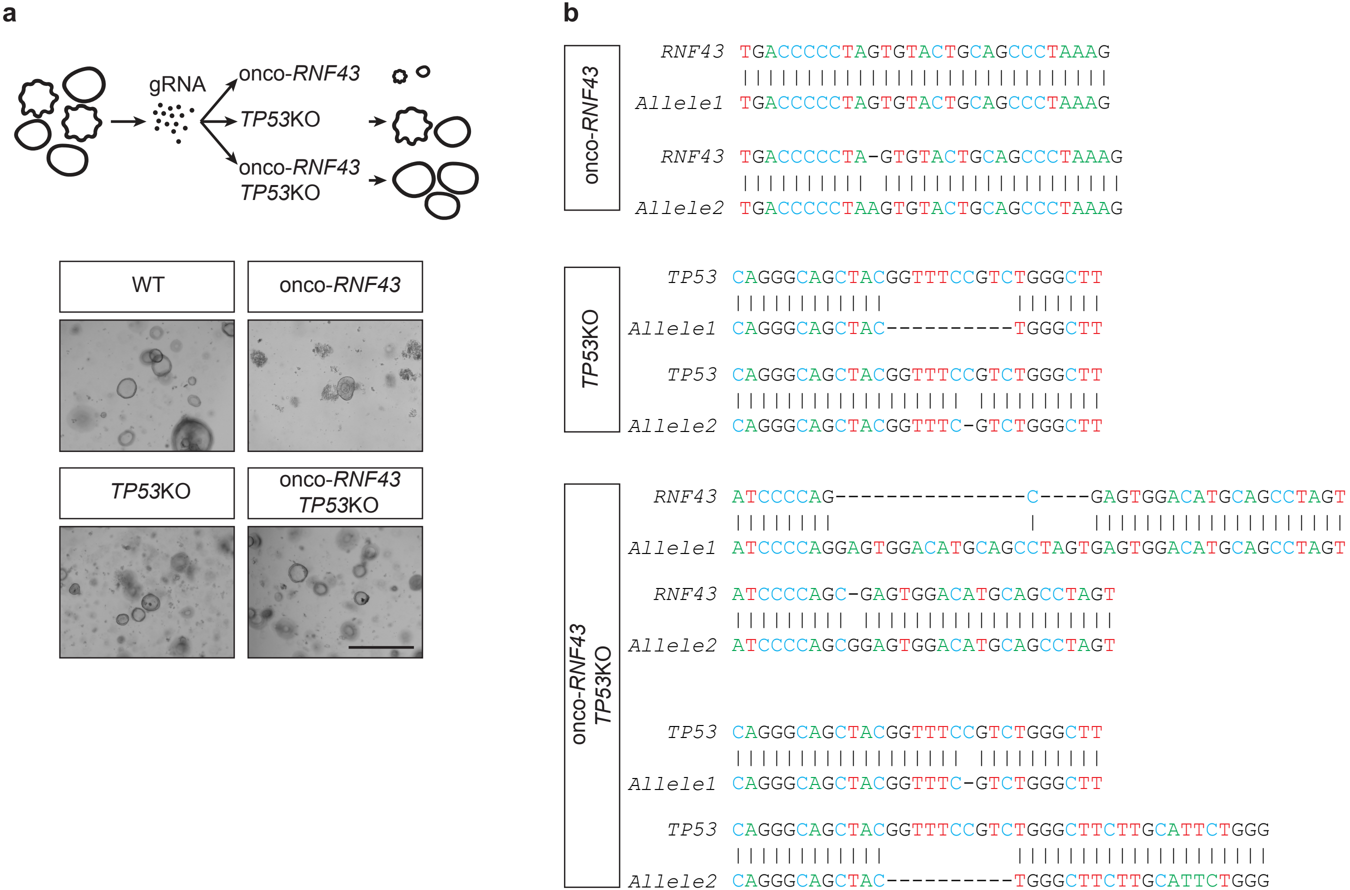

### Supplementary Figure 4

#### Supplementary Figure 4

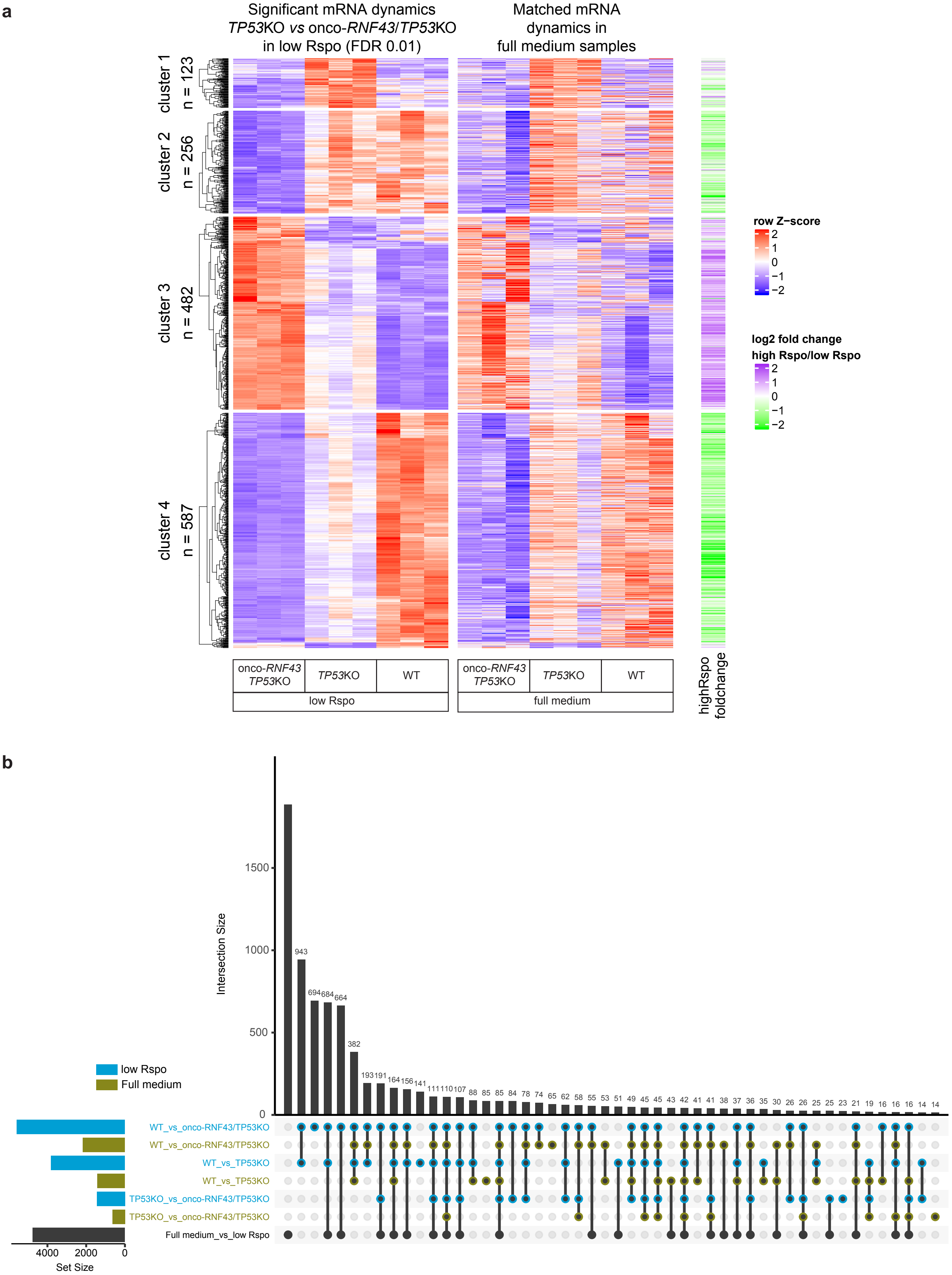
