## Supplementary Figure 5 for "RNF43 truncating mutations mediate a tumour suppressor-to-oncogene switch to drive niche-independent self-renewal in cancer"

### Supplementary blots

#### Figure 1b

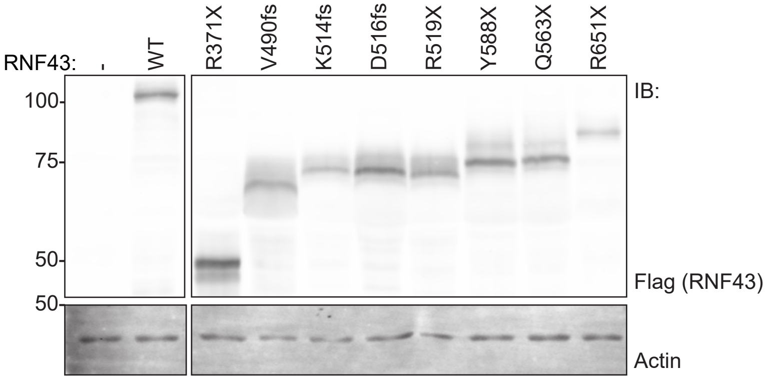


#### Figure 1g

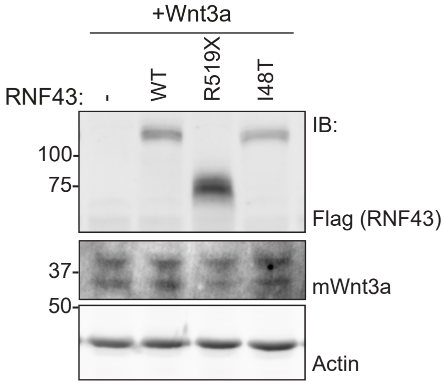


#### Figure 2b

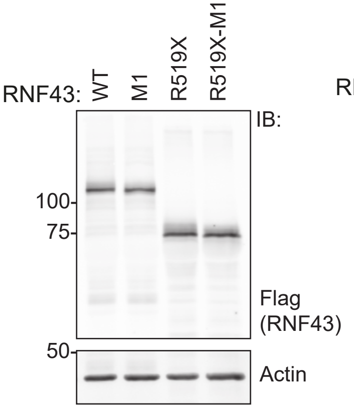


#### Figure 2c

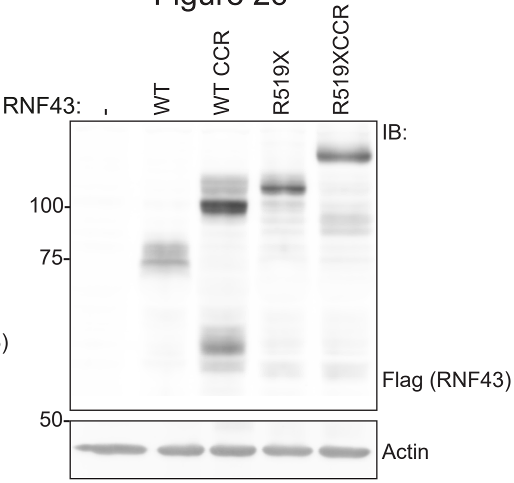


#### Figure 2d

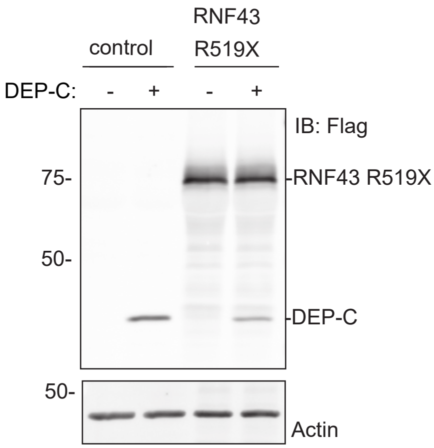


#### Figure 2e

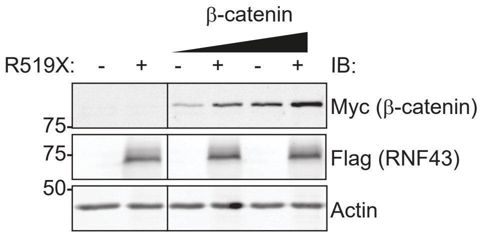


#### Figure 3e

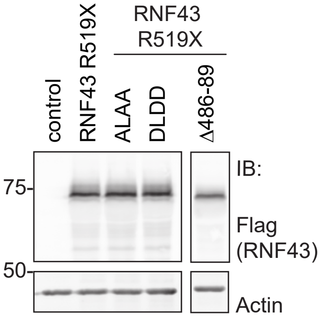


#### Supplementary Figure 1a

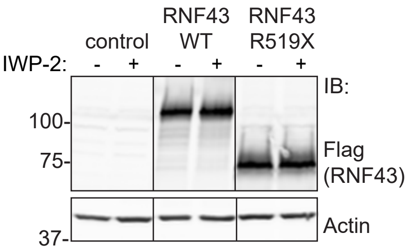


#### Supplementary Fig 1b

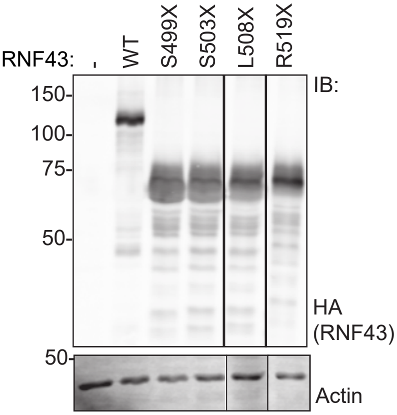


#### Supplementary Figure 2h

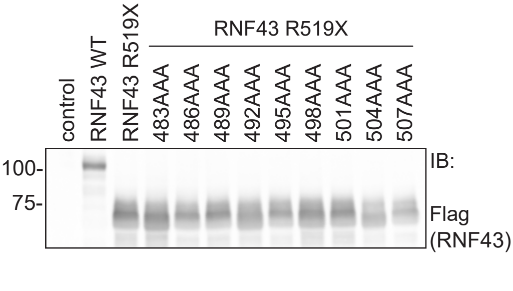


#### Supplementary Figure 3i

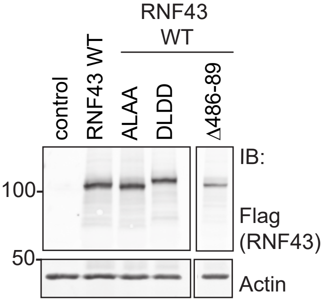
