## Supplementary Table 1 for "RNF43 truncating mutations mediate a tumour suppressor-to-oncogene switch to drive niche-independent self-renewal in cancer"

**Supplementary Table 1 | Onco-RNF43 mutations in human tumours.**

| Study | Sample ID | Cancer Type | Amino acid change | Mutations per sample | Mutations in senescence genes |
|---|---|---|---|---|---|
| Breast Invasive Carcinoma (TCGA, Nature 2012) | TCGA-AN-A04D-01 | Invasive Breast Carcinoma | K514Sfs*9 | 71 | RB1 (deep deletion) |
| TCGA data for Esophagus-Stomach Cancers (TCGA, Nature 2017) | TCGA-BR-8081-01 | Stomach Adenocarcinoma | D516Gfs*10 | 946 | TP53, ATM |
| Uterine Corpus Endometrial Carcinoma (TCGA, PanCancer Atlas) | TCGA-D1-A2G0-01 | Uterine Mixed Endometrial Carcinoma | D516Ifs*11 | 817 | PTEN |
| Merged Cohort of LGG and GBM (TCGA, Cell 2016) | TCGA-HT-7690-01 | Diffuse Glioma | R519* | 15 | TP53 |
| Uterine Corpus Endometrial Carcinoma (TCGA, Provisional) | TCGA-AP-A059-01 | Uterine Endometrioid Carcinoma | R519* | 9247 | TP53, ATM, ATR, RBL1, PTEN |
| Uterine Corpus Endometrial Carcinoma (TCGA, PanCancer Atlas) | TCGA-AX-A2HD-01 | Uterine Endometrioid Carcinoma | R519* | 7855 | ATM, TP63, RB1, RBL1, ATR, PTEN |
| Uterine Corpus Endometrial Carcinoma (TCGA, PanCancer Atlas) | TCGA-EY-A549-01 | Uterine Endometrioid Carcinoma | R519* | 1321 | TP63, ATR, RB1, PTEN |
| Colorectal Adenocarcinoma (DFCI, Cell Reports 2016) | coadread_dfci_2016_2945 | Colorectal Adenocarcinoma | S525Lfs*172 | 131 | TP53 |
| MSK-IMPACT Clinical Sequencing Cohort (MSKCC, Nat Med 2017) | P-0007969-T01-IM5 | Stomach Adenocarcinoma | T542Pfs*158 | 46 | ATR |
| Prostate Adenocarcinoma (TCGA, Provisional) | TCGA-CH-5746-01 | Prostate Adenocarcinoma | S546Qfs*50 | 32 | RBL1 (deep deletion) |
| Pan-Lung Cancer (TCGA, Nat Genet 2016) | TCGA-L9-A7SV-01 | Lung Adenocarcinoma | Y558* | 1341 | TP53 |
| MSK-IMPACT Clinical Sequencing Cohort (MSKCC, Nat Med 2017) | P-0012041-T01-IM5 | Pancreatic Adenocarcinoma | F562Sfs*138 | 8 | TP53, ATM, RB1 |
| Skin Cutaneous Melanoma (TCGA, Provisional) | TCGA-EE-A2GJ-06 | Cutaneous Melanoma | Q563* | 928 | TP53 |
