## Supplementary Table 2 for "RNF43 truncating mutations mediate a tumour suppressor-to-oncogene switch to drive niche-independent self-renewal in cancer"

**Supplementary Table 2 | Wnt destruction complex-associated proteins identified by BioID.**

| ID | Name | PD 1 | CTRL 1 | PD 2 | CTRL 2 | PD 3 | CTRL 3 |
|---|---|---|---|---|---|---|---|
| Q68DV7 | E3 ubiquitin-protein ligase RNF43 OS=Homo sapiens GN=RNF43 PE=1 SV=1 - [RNF43_HUMAN] | 68(68) | - | 69(69) | - | 73(73) | - |
| O14641 | Segment polarity protein dishevelled homolog DVL-2 OS=Homo sapiens GN=DVL2 PE=1 SV=1 - [DVL2_HUMAN] | 17(18) | - | 26(27) | - | 14(14) | - |
| Q92997 | Segment polarity protein dishevelled homolog DVL-3 OS=Homo sapiens GN=DVL3 PE=1 SV=2 - [DVL3_HUMAN] | 15(17) | - | 17(19) | - | 10(11) | - |
| O14640 | Segment polarity protein dishevelled homolog DVL-1 OS=Homo sapiens GN=DVL1 PE=1 SV=2 - [DVL1_HUMAN] | 12(13) | - | 16(17) | - | 14(15) | - |
| P25054 | Adenomatous polyposis coli protein OS=Homo sapiens GN=APC PE=1 SV=2 - [APC_HUMAN] | 26(26) | - | 22(22) | - | 17(17) | - |
| O15169 | Axin-1 OS=Homo sapiens GN=AXIN1 PE=1 SV=2 - [AXIN1_HUMAN] | 5(5) | - | 9(9) | - | 5(5) | - |
| P49674 | Casein kinase I isoform epsilon OS=Homo sapiens GN=CSNK1E PE=1 SV=1 - [CK1E_HUMAN] | 2(2) | - | 3(4) | - | 2(2) | - |
| P48729 | Casein kinase I isoform alpha OS=Homo sapiens GN=CSNK1A1 PE=1 SV=2 - [CK1A_HUMAN] | 2(2) | - | 2(2) | - | 2(2) | - |

Number of identified unique peptides are shown. Total identified peptides are indicated between brackets. PD, pull down; CTRL, control.
