## Supplementary Table 3 for "RNF43 truncating mutations mediate a tumour suppressor-to-oncogene switch to drive niche-independent self-renewal in cancer"

**Supplementary Table 3 | Phosphopeptides identified in RNF43 WT and R519X.**

| Phosphosites | # P | RNF43 WT | | | RNF43 R519X | | |
|---|---|---|---|---|---|---|---|
| | | no kinase | Ck1α | Ck1ε | no kinase | Ck1α | Ck1ε |
| S241 | 1 | ND | ND | ND | 7.37 | 7.20 | 7.21 |
| S241-T245 | 1 | ND | ND | ND | 6.67 | 6.44 | 6.76 |
| T308 | 1 | ND | ND | ND | 6.93 | 7.07 | 7.21 |
| T317 | 1 | ND | ND | ND | 6.80 | 7.04 | 7.16 |
| S321 | 1 | ND | ND | ND | 8.54 | 8.30 | 8.60 |
| S325 | 1 | 6.65 | 5.85 | 6.32 | 7.23 | 7.10 | 7.40 |
| S323-S325 | 1 | ND | ND | ND | 7.49 | 7.48 | 7.82 |
| S331 | 1 | 5.74 | 6.26 | 5.40 | 7.37 | 7.33 | 7.62 |
| S443-S446 | 2 | 6.48 | 7.45 | 7.12 | 8.45 | 8.36 | 9.05 |
| S446 | 1 | 7.31 | 7.64 | 7.49 | 8.61 | 8.61 | 9.03 |
| S443, S444, S446 | 3 | NQ | NQ | NQ | ND | NQ | ND |
| S499-S503 | 1 | ND | ND | ND | 7.00 | 7.08 | 7.58 |
| S499-S503 | 2 | ND | ND | ND | ND | 6.14 | 6.28 |
| S512 | 1 | 6.79 | 7.03 | 6.43 | 7.49 | 7.09 | 7.52 |
| S532 | 1 | 8.52 | 8.63 | 8.33 | ND | ND | ND |
| S535 | 1 | 6.47 | 6.83 | 6.31 | ND | ND | ND |
| S532, S535 | 2 | ND | 7.37 | ND | ND | ND | ND |
| S525-S535, T539 | 2 | ND | 6.74 | ND | ND | ND | ND |
| S607-S611 | 1 | 7.50 | 7.85 | 7.13 | ND | ND | ND |
| S611 | 1 | 6.19 | 6.60 | 6.13 | ND | ND | ND |

The table summarises the intensities of the identified phosphopeptides in the $Log_{10}$ scale. Areas of indicated phosphopeptides were normalised. Phosphopeptide standards added prior to $TiO_2$ fractionation were used for normalisation. Non-phosphorylated peptides were used for the second part of normalisation. Individual areas of the phosphopeptides were combined and presented as the total area of each phosphosite.

| | |
|---|---|
| Phosphosite: | position of phosphosite in the RNF43 protein sequence phosphosites |
| separated by a dash: | phosphorylation was detected and occurs on one (or more) amino acids within the listed range, but position cannot be determined with certainty |
| # P: | number of phosphorylations observed on the peptide for the specific peptide spectrum match (PSM) |
| ND: | not detected |
| NQ: | identified, but cannot be quantified |
| In grey: | below detection limit, for quality only |
